# Genetic ancestry predicts male-female affiliation in a natural baboon hybrid zone

**DOI:** 10.1101/2020.10.28.358002

**Authors:** Arielle S. Fogel, Emily M. McLean, Jacob B. Gordon, Elizabeth A. Archie, Jenny Tung, Susan C. Alberts

## Abstract

Opposite-sex social relationships are important predictors of fitness in many animals, including several group-living mammals. Consequently, understanding sources of variance in the tendency to form opposite-sex relationships is important for understanding social evolution. Genetic contributions are of particular interest due to their importance in long-term evolutionary change, but little is known about genetic effects on male-female relationships in social mammals, especially outside of the mating context. Here, we investigate the effects of genetic ancestry on male-female affiliative behavior in a hybrid zone between the yellow baboon (*Papio cynocephalus*) and the anubis baboon (*P. anubis*), in a population in which male-female social bonds are known predictors of lifespan. We place our analysis within the context of other social and demographic predictors of affiliative behavior in baboons. Genetic ancestry was the most consistent predictor of opposite-sex affiliative behavior we observed, with the exception of strong effects of dominance rank. Our results show that increased anubis genetic ancestry is associated with subtly, but significantly higher rates of opposite-sex affiliative behavior, in both males and females. Additionally, pairs of anubis-like males and anubis-like females were the most likely to socially affiliate, resulting in moderate assortativity in grooming and proximity behavior as a function of genetic ancestry. Our findings indicate that opposite-sex affiliative behavior partially diverged during baboon evolution to differentiate yellow and anubis baboons, despite overall similarities in their social structures and mating systems. Further, they suggest that affiliative behavior may simultaneously promote and constrain baboon admixture, through additive and assortative effects of ancestry, respectively.

**HIGHLIGHTS:**

- Opposite-sex social relationships can have important fitness consequences.
- In hybrid baboons, genetic ancestry predicted male-female affiliative behavior.
- Both an individual’s genetic ancestry and that of its social partner mattered.
- Male-female affiliation was assortative with respect to genetic ancestry.
- Dominance rank and group demography also influenced male-female social affiliation.

---

Social relationships, both within and between sexes, are ubiquitous features in the lives of social mammals. Affiliative interactions among members of the same sex are positively associated with fertility or survival in a number of social mammal species, including group-living primates, equids, cetaceans, and rodents (e.g., Cameron et al., 2009; Ellis et al., 2019; Frère et al., 2010a; Schülke et al., 2010; Silk et al., 2009, 2010; Weidt et al., 2008). Opposite-sex affiliative bonds can also have important consequences. In monogamous species, strong social bonds between sexual partners predict shorter interbirth intervals, increased offspring number, and improved offspring survival, potentially due to improved coordination between partners in caring for young, obtaining resources, or defense against predators (e.g., Black, 2001; Griggio & Hoi, 2011; Ribble, 1992; Sánchez-Macouzet et al., 2014). Further, in some group-living primates, females compete for access to males outside of mating contexts, suggesting that social bonds with males are themselves an important resource (Archie et al., 2014; Baniel et al., 2016, 2018; Cheney et al., 2012; Haunhorst et al., 2019; Lemasson et al., 2008; Palombit et al., 2001; Seyfarth, 1978). In support of this idea, females in several cercopithecine monkey species benefit from opposite-sex social bonds via enhanced survival, care for their offspring, and protection from harassment (Archie et al., 2014; Baniel et al., 2016; Haunhorst et al., 2017; Kulik et al., 2012; Lemasson et al., 2008; Moscovice et al., 2009; Nguyen et al., 2009; Palombit et al., 1997; Seyfarth, 1978; Silk et al., 2020; Weingrill, 2000). Males of these species may also benefit from social bonds with females. For example, baboon males who form strong social bonds with females tend to live longer than those who do not (Campos et al., 2020). Males may also benefit by gaining mating opportunities (although the evidence for this benefit is mixed), opportunities to care for their offspring, or access to infants that can be exploited for social gain (Ménard et al., 2001; Packer, 1979b; Smuts, 1985; van Schaik & Paul, 1996; Whitten, 1987).

While a number of studies have investigated the sources of variance in same-sex affiliative relationships in group-living mammals (Best et al., 2014; Frère et al., 2010b; Langergraber et al., 2009; Mitani, 2009; Möller et al., 2001; Seyfarth, 1976; Seyfarth et al., 2014; Silk et al., 2006a; Silk et al., 2006b; Smith et al., 2006; Widdig et al., 2001), we know comparably less about the sources of variance in opposite-sex relationships, especially outside the mating context. Addressing this gap is important for understanding the evolution of heterosexual bonds. In particular, if the tendency to form opposite-sex social bonds is affected by genotype, it has the potential to evolve in response to natural selection. Strong evidence for genetic effects comes from interspecific comparisons between pair-bonded and multiply mating species. For example, comparisons between the monogamous prairie vole and other, closely related promiscuous voles have identified genetic divergence in the pathways that regulate arginine vasopressin, oxytocin, and dopamine signaling, which in turn influences pair-bonding behavior (Young et al., 1996; Young et al., 1999; Young et al., 1997a; Young et al., 1997b; reviewed in Carter & Perkeybile, 2018; Johnson & Young, 2015; Sadino & Donaldson, 2018; Young et al., 2011). These pathway differences may in part be due to differences in the distribution and densities of hormone receptors in the brain, suggesting one important mechanism through which variation in opposite-sex social relationships evolves (Insel & Shapiro, 1992; Insel et al., 1994; Smeltzer et al., 2006). Research in other pair-bonded rodents, primates, fish, frogs, and birds has placed these findings in a broader context, indicating that these and other pathways (e.g., Young et al., 2019) consistently influence pair-bonding across divergent species, although they may do so in a species-specific manner (reviewed in Carter & Perkeybile, 2018; Fischer et al., 2019a; Johnson & Young, 2015).

Despite these important discoveries in pair-bonded species, little is known about genetic influences on opposite-sex social bonding in group-living animals, including the degree to which genotype contributes to differences between species with similar social and mating systems. Here, we investigate the association between genetic ancestry and male-female affiliative behavior in a well-studied natural primate population, the baboons of Kenya’s Amboseli basin (Alberts, 2018; Alberts & Altmann, 2012). Baboons (genus *Papio*) began speciating ~1.4 million years ago, and today, the six extant species occupy distinct geographic ranges across Africa (Rogers et al., 2019). Most species of baboons, including those in Amboseli, live in multi-male, multi-female social groups in which multiple individuals of both sexes mate and form social bonds (Fischer et al., 2019b). Amboseli lies in a hybrid zone between two such species, the yellow baboon (*Papio cynocephalus*) and the anubis baboon (*P. anubis*, also known as the olive baboon) (Alberts & Altmann, 2001; Samuels & Altmann, 1986; Tung et al., 2008; Wall et al., 2016). While yellow baboons contribute the majority of genetic ancestry in this population, the range of admixture we observe—from animals that are almost entirely yellow to those that are almost entirely anubis—gives us the opportunity to examine potential genetic ancestry effects on opposite-sex affiliative relationships. Complementary data on social and demographic variables for the same individuals allow us to place these effects in the context of other, environmental sources of variance.

## Genetic ancestry effects on male-female interactions in hybrid zones

The Amboseli baboon hybrid zone provides a “natural laboratory” for understanding the relationship between genetic ancestry and affiliative behavior because it allows individuals with varyingly admixed genomes to be observed in a shared environment (Hewitt, 1988). In turn, studying social behavior in hybrid zones can shed light onto hybrid zone dynamics, as most clearly illustrated in cases where ancestry influences mating behavior. In such cases, assortative mating by ancestry limits gene flow and can reinforce species boundaries, whereas ancestry-related mating advantages can lead to asymmetric gene flow and range expansion (e.g., Baldassarre & Webster, 2013; Baldassarre et al., 2014; Kronforst et al., 2006; Mavárez et al., 2006).

Ancestry effects on mating behavior have also been detected in both the yellow baboon-anubis baboon hybrid zone in Amboseli and in an anubis baboon-hamadryas baboon hybrid zone in Ethiopia. In Amboseli, anubis-like males are more likely to obtain consortships (extended mate-guarding associations between an adult male and an adult female in estrus, during which most conceptions occur), and male-female pairs with similar genetic ancestry are more likely to consort than pairs with different ancestry (Tung et al., 2012). In the Ethiopian hybrid zone, ancestry affects both male mating strategy and how females respond to males (Bergman & Beehner, 2003). However, we do not yet understand whether genetic ancestry effects extend to other aspects of male-female interactions, such as affiliation between male-female pairs outside of the mating context (but see Bergman et al., 2008 for an analysis of male interest in non-estrus females). If so, genetic ancestry effects on male-female social relationships may be more important than indicated by analyses of mating behavior alone. Specifically, because opposite-sex social affiliation also predicts lifespan in the Amboseli population (Archie et al., 2014; Campos et al., 2020), ancestry effects on this trait may secondarily affect how long individuals live and who they co-reside with, thus influencing the genetic composition of subsequent generations.

## Goals of this study

Here, we evaluated the extent to which genetic ancestry predicts the formation of male-female social relationships in baboons. We focused specifically on male-female affiliative behavior in non-mating contexts (i.e., periods when females were pregnant or lactating, and not sexually cycling) because social relationships in these contexts are not driven by immediate sexual interactions. Using two multivariate models (one for grooming behavior and one for proximity behavior), we simultaneously tested for (i) the additive effects of male and female individual characteristics, including genetic ancestry, on the probability of affiliative social behavior between males and females, and (ii) characteristics defined by the pair, including ancestry-related assortativity. In the same model, we also tested two additional hypotheses: (iii) that opposite-sex affiliation depends on female reproductive state (i.e., pregnancy or lactation), based on evidence that the stability of male-female relationships varies across baboon species as a function of female reproductive state (Baniel et al., 2016; Fischer et al., 2017; Goffe et al., 2016; Nguyen et al., 2009; Städele et al., 2019; Weingrill, 2000); and (iv) that opposite-sex affiliation depends on group demography, based on findings that male-female interactions in baboons and other primates also depend on group composition (Archie et al., 2014; Bergman & Beehner, 2003; Rosenbaum et al., 2016; Tung et al., 2012).

## METHODS

### Study subjects

Study subjects were adult baboons from an intensively studied wild population inhabiting the Amboseli ecosystem of southern Kenya (Alberts, 2018; Alberts & Altmann, 2012). This population consists of multigeneration hybrids, most of which have predominantly yellow baboon ancestry, but some of which are recent hybrid descendants of anubis or anubis-like immigrants that have arrived in Amboseli since the early 1980’s (approximately a decade after long-term observations began) (Alberts & Altmann, 2001; Charpentier et al., 2008; Samuels & Altmann, 1986; Tung et al., 2008; Wall et al., 2016). This natural hybrid population is situated within a narrow hybrid zone that likely extends along the geographic boundary between yellow baboon and anubis baboon distributions in East Africa (Charpentier et al., 2012).

Members of the Amboseli baboon study population are individually recognized based on physical appearance and are monitored on a near-daily basis by trained observers who record demographic data (e.g., group membership, births, deaths, immigration, emigration) and behavioral data (e.g., social interactions, mating, traveling, resting, feeding). Study subjects were parous adult females (because parous females are strongly preferred over nulliparous females as mates by adult males of most primate species: Anderson, 1986; Gesquiere et al., 2007) and adult males that had achieved a social dominance rank among other adult males in their group (Table S1). Overall, we considered members of twelve social groups that were studied between November 1999 and December 2015. We restricted the data set to include only males and females for whom estimates of genetic ancestry, genetic diversity, and genetic relatedness between individuals in male-female pairs could be calculated from previously generated microsatellite data (Buchan et al., 2003; Tung et al., 2008; Tung et al., 2012). The resulting sample contained 136 females and 160 males, who together formed 3,468 unique male-female dyads across the grooming and proximity data sets.

### Affiliative social behavior

Grooming and maintenance of close spatial proximity (hereafter, proximity) are affiliative behaviors important to establishing, maintaining, and strengthening social bonds in non-human primates (Cords, 1997; Cords, 2012; Palombit et al., 1997; Silk et al., 2013). Although male-female grooming and proximity events were moderately correlated in our data set (Pearson’s product-moment correlation: *r* = 0.222, *P* < 10^−15^), we analyzed grooming and proximity separately because grooming measures only explained 4.9% of the variance in the proximity data. Grooming data were collected during systematic monitoring of the population, following a sampling protocol in which observers move in a predetermined random order throughout the group. This approach avoids biases due to uneven sampling of subjects (Alberts et al., 2020; Archie et al., 2014). Proximity data were collected during random-order focal animal sampling on adult females, during which the identity of the nearest adult male within 5 meters, if any, was recorded once per minute for the duration of each 10 minute sample (Alberts et al., 2020). We excluded data from time periods in which behavioral monitoring was inconsistent or when social groups were too unstable (i.e., social groups were fissioning or fusing) to unambiguously determine an individual’s group membership. We also excluded all data from the 2009 hydrological year (November 1^st^, 2008-October 31^st^, 2009) which included the most severe drought documented in the Amboseli basin in more than 40 years (Okello et al., 2016; Tuqa et al., 2014). Omitting data from 2009 ensured that effects from this rare and extreme event, which altered patterns of female fertility and reproductive states, did not influence our results (Fitzpatrick et al., 2014; Lea et al., 2015).

### Predictor variables

We investigated the relationship between genetic ancestry and opposite-sex affiliative social behavior using the following predictors, motivated in part by known predictors of mating behavior in this population (Tung et al., 2012) (see Tables S2-S3 for correlations among all predictor variables).

#### Genetic ancestry

Genetic estimates of hybridity (i.e., the proportion of each individual’s genome estimated to be from anubis ancestry) were included for females (*h_f_*) and males (*h_m_*). These estimates were based on genotypes at up to 13 highly polymorphic microsatellite markers and average ancestry assignments produced using the Bayesian clustering algorithm STRUCTURE 2.3.4 (Falush et al., 2003; Pritchard et al., 2000; see Tung et al., 2008; mean typed loci per individual = 12.40 ± 1.10 s.d.). These assignments range continuously from 0 to 1, where 0 corresponds to unadmixed yellow baboon ancestry and 1 corresponds to unadmixed anubis baboon ancestry. These estimates are strongly correlated with recent genome-wide ancestry estimates (Pearson’s product-moment correlation: *r* = 0.717, *P* = 1.17 × 10^−4^, n=23 individuals that overlapped between data sets) (Wall et al., 2016); however, because genome-wide estimates are available for only a subset of the population, we used the microsatellite-based estimates here.

#### Assortative genetic ancestry index

To test the possibility that males and females of similar genetic ancestry are more likely to socially affiliate, we calculated a pairwise assortative genetic ancestry index, *b*, as a function of the genetic ancestry estimates of the female and male (*h_f_* and *h_m_*, respectively), paralleling the approach used in Tung et al. (2012)’s pairwise assortative mating index, *a*:

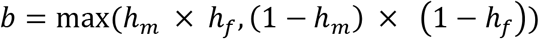

This index ranges from 0 to 1: high values indicate highly assortative male-female pairs (i.e., individuals in the pair both have low or high genetic ancestry estimates) and low values indicate highly disassortative male-female pairs (i.e., individuals in the pair have different genetic ancestry estimates). Intermediate values indicate male-female pairs where both individuals are of intermediate ancestry.

#### Heterozygosity

High genetic diversity is sometimes thought to be a measure of genetic quality (Kempenaers, 2007). Because it is relevant to mate choice (Kempenaers, 2007) and potentially social partner choice, we therefore included a measure of genetic diversity for both males and females using up to 14 highly polymorphic microsatellite markers (mean typed loci per individual = 13.13 ± 1.22 s.d.; 13 of these markers were also used to assign genetic ancestry scores). We estimated individual genetic diversity by dividing the number of heterozygous loci by the number of genotyped loci for each individual (following Charpentier et al., 2008). Importantly, there is no overall effect of species identity (i.e., yellow or anubis) on genetic diversity using these markers (Charpentier et al., 2012).

#### Relatedness

Because the formation of social bonds may be affected by kinship, we included an estimate of genetic relatedness for each male-female dyad using the method of Queller and Goodnight (1989). These estimates, based on the same genotype data used to estimate heterozygosity, were calculated using the function *coancestry* in the R package *related* (version 1.0; Pew et al., 2015; Wang, 2011).

#### Social dominance rank

Social dominance rank can enhance access to valuable resources, including desirable social partners (e.g., Archie et al., 2014; Baniel et al., 2016; Haunhorst et al., 2019; Lemasson et al., 2008; Palombit et al., 2001). We therefore modeled female rank, male rank, and the interaction between female and male ranks as additional fixed effects in the models. Female and male ranks were assigned separately for each sex, on a monthly basis, based on the outcomes of dyadic agonisms between all pairs of individuals in the same group (Alberts et al., 2020). We represented rank using an ordinal approach, where the highest-ranking individual holds rank 1 and lower-ranking individuals occupy ranks of successively higher numbers. Since female and male ranks were assigned on a monthly basis and our time window for analyses of grooming and proximity interactions spanned a two-month period, we used the average of each individual’s rank across both months for each two-month interval.

#### Age

Female age may also affect a female’s social interactions. To account for possible age-related effects, we modeled a linear effect of female age, averaged across each two-month analysis window (i.e., her age at the start of the second month), as a continuous predictor variable in our models. We also included a transformed measure of female age that reflects the relationship between female age and conception probability in this population, where the highest conception probability occurs at ~14 years of age (Beehner et al., 2006). Following Tung et al. (2012), we calculated female transformed age, *a_t_*, as a function of *a_u_*, the untransformed female age:

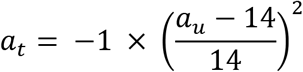

This transformation assigns 0 for the value of *a_t_* at 14, the age at which conception probabilities are highest; values of *a_t_* become increasingly negative with distance from age 14. For 90.4% (123 out of 136) of the females in the data set, birthdates were known to within a few days. For the other females in the data set, birthdates were estimated to within 6 months (i.e., ± 3 months’ error). Male age was not included in any models since it is tightly correlated with rank in male baboons (Alberts et al., 2003) and its effect on mating and social behavior is likely to be linked to rank (Silk et al., 2020; Tung et al., 2012).

#### Group composition

To incorporate group-level demographic effects on social behavior, we included the number of adult females and the number of adult males in the social group of a male-female pair in both models (averaged across each two-month analysis period).

#### Reproductive state

Because female reproductive state affects the stability of male-female bonds in other baboon species (Baniel et al., 2016; Weingrill, 2000), we also included female reproductive state as a categorical variable in our models. To capture opposite-sex affiliation outside the context of mating, we excluded all data points in which the female member of a potential pair was cycling. Thus, reproductive state was either pregnant or lactating, both of which meant that the female was not actively mating and could not conceive. Pregnancy and lactation were coded as −1 and 1, respectively, which avoided numerical instability that occurred if we used a 0/1 encoding (see Supplementary Methods). To test whether the effects of female reproductive state on male-female social affiliation depended on genetic ancestry, we also modeled an interaction between female reproductive state and female genetic ancestry and a separate interaction between female reproductive state and male genetic ancestry.

#### Pair co-residency

The number of days that a male and female were observed in the same social group may influence both their tendency to affiliate and our ability to detect interactions between them. We therefore included the total number of days in each two-month interval that a male and female were censused in the same group as a model covariate.

#### Observer effort

The number of field observers and the amount of time spent conducting behavioral observations for each study group was consistent across all study groups regardless of their size (Fig. S1). Consequently, the probability of observing grooming or proximity events could vary as a function of social group size, because an observer watching a small group is likely to capture a larger fraction of interactions in a given time period than that same observer watching a much larger group. Thus, we calculated observer effort and included it as a covariate in both models. Observer effort was estimated as the average number of minutes of focal sample data collected per adult female per social group in a given two-month interval (see Supplementary Methods).

### Statistical analyses

Grooming and proximity behavior were modeled as binary events and analyzed separately using binomial mixed effects models. Each row of data corresponded to a unique, co-resident female-male dyad in a given two-month interval, and was assigned a value of “1” if the dyad was observed grooming or in proximity at least once during the two-month interval and a “0” if they were not. We used two months as our time interval because our resolution for grooming and proximity behavior is relatively coarse on a month-to-month basis, even after excluding months in which observer effort was low (see Supplementary Methods).

We retained all two-month intervals in which focal females groomed or were in proximity with any candidate male social partner at least once, except for: (i) two-month intervals in which females transitioned between reproductive states; and (ii) two-month intervals in which the average number of adult males in the social group was less than two. We also excluded any male in a female’s two-month interval if he was only present for one of the two months and excluded all data for females and males who were observed for less than 8 months because sparse data makes it difficult to estimate individual-level random effects. The final grooming data set included 127 unique females and 160 unique males across 1,866 female two-month intervals (17,356 female-male pair-interval combinations), and the final proximity data set included 131 unique females and 160 unique males across 2,338 female two-month intervals (21,130 total female-male pair-interval combinations).

We ran binomial mixed effects models using the function *glmmTMB* (family = “binomial”) in the R package *glmmTMB* (version 1.0.1; Brooks et al., 2017), using a logit link:

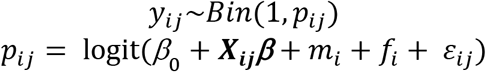

where *y_ij_* is a 0/1 value indicating whether male-female dyad *i* was observed grooming or in proximity during a two-month interval *j*. *y_ij_* is drawn from a binomial distribution, where the probability of grooming or proximity (*p_ij_*) is modeled as the function of the logit-transformed sum of (i) the intercept, *β*_0_; (ii) the fixed effects (***X_ij_β***) of male genetic ancestry, female genetic ancestry, the assortative genetic ancestry index for that pair, male heterozygosity, female heterozygosity, genetic relatedness between individuals in that pair, male dominance rank, female dominance rank, female age, transformed female age, the number of adult females in the social group, the number of adult males in the social group, female reproductive state (pregnant or lactating), the interaction between female reproductive state and female genetic ancestry, the interaction between female reproductive state and male genetic ancestry, the interaction between male and female dominance ranks, pair co-residency, and observer effort (***X_ij_*** represents all of these data using standard matrix notation and ***β*** refers to the vector of all fixed effect estimates); and (iii) the random effects of male identity, *m_i_*, and female identity, *f_i_*. *ε_ij_* represents model error.

To assess statistical significance, we used permutation tests to account for unequal representation of individuals in the data set and predictors that did not follow standard parametric distributions. We followed the procedure of Tung et al. (2012), who conducted a similar analysis on mating behavior. Specifically, we first computed, for each female-interval combination, the proportion of dyads where an event (grooming or proximity) occurred. These values are estimates of the probability of grooming or proximity with any male, per female-interval combination. These probabilities were then permuted across all female two-month intervals, and randomized response variables (0/1) were generated by drawing from a binomial distribution with *P_ij_* equal to the permuted grooming or proximity probability for each female-interval. This approach preserves the structure of the predictor variables (including correlations between predictors), the number of times each individual is represented in the data set, and the distribution of grooming or proximity events for each female-interval. We then fit the model used to analyze the real data to the permuted data set and calculated a p-value for each predictor variable based on the number of times that the absolute value of the effect size estimated from the permuted data sets was greater than the absolute value of the effect size estimated from the observed data set, across 1,000 permutations. All analyses were run in R (version 3.6.1; R Core Team, 2019).

### Ethical note

The research in this study was approved by the Institutional Animal Care and Use Committees (IACUC) at Duke University (#A273-17-12), and adhered to the laws and guidelines of the Kenyan government.

## RESULTS

### Individual characteristics: genetic ancestry and dominance rank predict opposite-sex affiliative social behavior in males and females

Our models identified two male characteristics that consistently predicted opposite-sex grooming and proximity behavior (Tables 1–2). Specifically, grooming and proximity were more likely to occur if the male in the dyad had more anubis ancestry (grooming: β = 0.429, p < 0.001, Table 1, Fig. 1a; proximity: β = 0.270, p < 0.001, Table 2, Fig. S2a) and was higher ranking (grooming: β = −0.096, p < 0.001, Table 1, Fig. 1b; proximity: β = −0.047, p < 0.001, Table 2, Fig. S2b). Male heterozygosity was not significantly associated with either grooming or proximity behavior.

**Figure 1.**
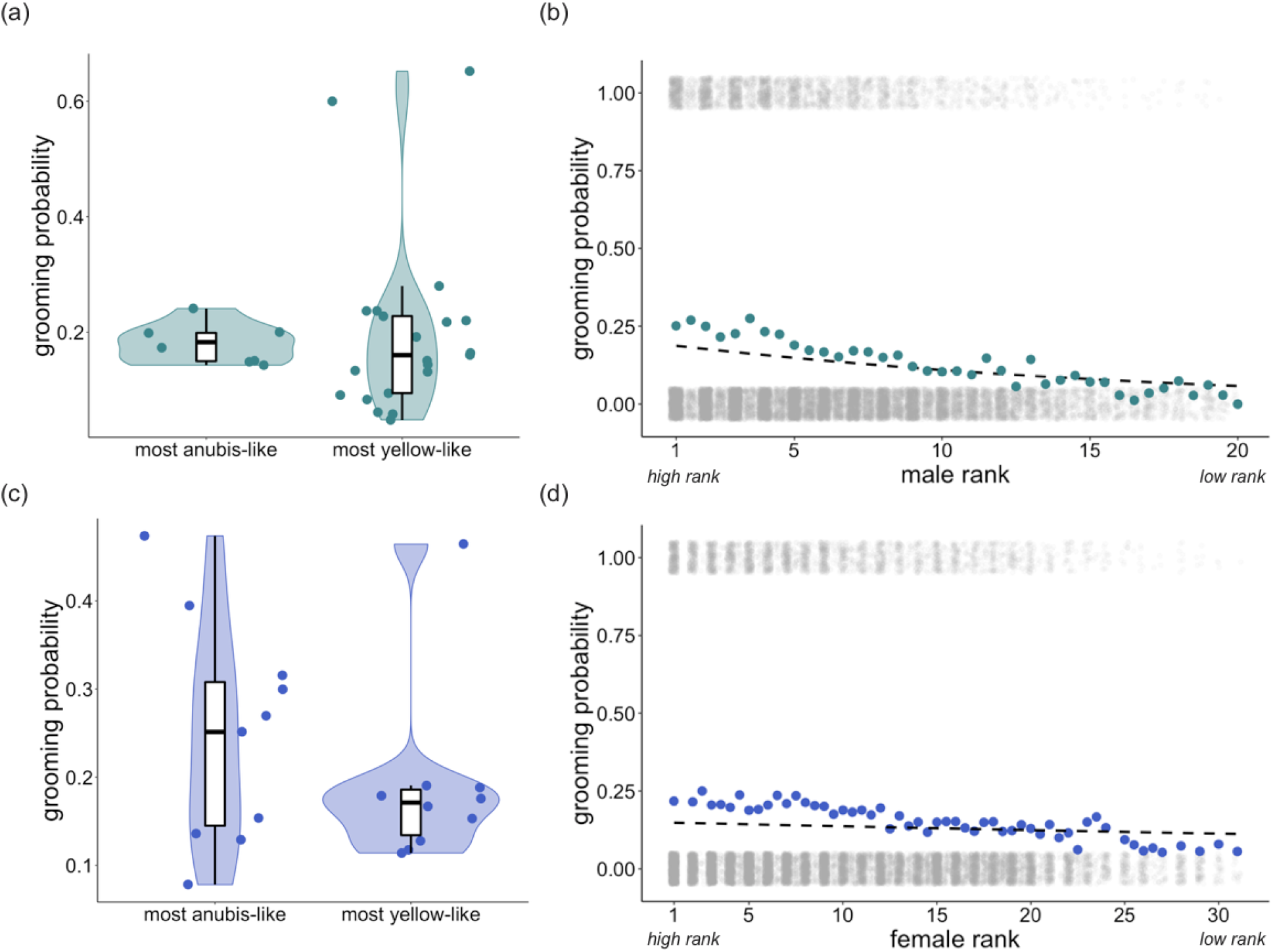
Genetic ancestry and dominance rank predict the tendency to groom with an opposite-sex partner. **(a)** The probability of grooming among co-resident opposite-sex pairs, per two-month interval, for the most anubis-like males (above the 90^th^ percentile for male genetic ancestry in the data set, > 83.6% anubis ancestry, n=8 males) and the most yellow-like males (below the 10^th^ percentile for male genetic ancestry in the data set, < 4.8% anubis ancestry, n=21 males). Probabilities were calculated from the data without adjustment for other covariates. **(b)** The probability of grooming among co-resident opposite-sex pairs, per two-month interval, as a function of male dominance rank. Colored dots show probabilities based on counts of grooming occurrences, without adjustment for other covariates (as in (a)), and the dashed line shows the predicted relationship based on model estimates, assuming average values for all other covariates (see Supplementary Methods). Grey dots show the presence (y=1) or absence (y=0) of grooming behavior for all 17,356 female-male pair-interval combinations, as a function of male dominance rank (dots are jittered vertically for visibility). Non-integer values correspond to individuals that changed ranks during a two-month interval in the data set. **(c**) As in (a), for the most anubis-like females (above the 90^th^ percentile for female genetic ancestry in the data set, > 76.0% anubis ancestry, n=11 females) and the most yellow-like females (below the 10^th^ percentile for female genetic ancestry in the data set, < 3.5% anubis ancestry, n=10 females). **(d)** As in (b), with the probability of grooming shown as a function of female dominance rank.

**Table 1.** Results from a multivariate logistic regression model predicting grooming behavior.

| Predictor variable<sup>a</sup> |  | Effect estimate | p-value<sup>b</sup> | Effect direction<sup>c</sup> |
| --- | --- | --- | --- | --- |
| Intercept |  | -1.474 | 0.339 | - |
| Genetic effects | **Female genetic ancestry** | **0.513** | **<0.001** | **more anubis ancestry in females → ↑ Pr(groom)** |
|  | **Male genetic ancestry** | **0.429** | **<0.001** | **more anubis ancestry in males → ↑ Pr(groom)** |
|  | **Assortative genetic ancestry index (*b*)** | **0.646** | **<0.001** | **females and males of similar genetic ancestry → ↑ Pr(groom)** |
|  | Female heterozygosity | -0.249 | 0.406 | - |
|  | Male heterozygosity | 0.228 | 0.189 | - |
|  | Genetic relatedness | -0.090 | 0.489 | - |
| Rank effects | **Female ordinal rank** | **-0.027** | **<0.001** | **higher ranking females → ↑ Pr(groom)** |
|  | **Male ordinal rank** | **-0.096** | **<0.001** | **higher ranking males → ↑ Pr(groom)** |
|  | **Female ordinal rank × male ordinal rank** | **0.003** | **<0.001** | **females and males of similar rank → ↑ Pr(groom)** |
| Age effects | Female age ($a_u$) | -0.007 | 0.332 | - |
| | Female age transformed ($a_t$) | -0.192 | 0.562 | - |
| Demographic effects | **Adult females in group** | **-0.039** | **<0.001** | **more adult females in group → ↓ Pr(groom)** |
|  | *Adult males in group* | *-0.030* | *0.018* | *more adult males in group → ↓ Pr(groom)* |
| Reproductive state effects | Reproductive state (pregnant = -1, lactating = +1) | -0.067 | 0.165 | - |
|  | Reproductive state (as above) × female genetic ancestry | 0.086 | 0.412 | - |
|  | Reproductive state (as above) × male genetic ancestry | 0.004 | 0.949 | - |
| Co-residency effects | **Pair co-residency** | **0.047** | **<0.001** | **longer co-residency → ↑ Pr(groom)** |
| Observer effects | Observer effort | 0.002 | 0.425 | - |

<sup>a</sup> All variables included in this table were fit as fixed effects in the multivariate logistic regression model. Male and female identity were fit as random effects.
<sup>b</sup> Predictor variables for which $p < 0.01$ are bolded and $p < 0.05$ are italicized.
<sup>c</sup> Pr(groom) = probability of grooming.

**Table 2.** Results from a multivariate logistic regression model predicting proximity behavior.

| Predictor variable<sup>a</sup> |  | Effect estimate | p-value<sup>b</sup> | Effect direction<sup>c</sup> |
| --- | --- | --- | --- | --- |
| Intercept |  | -1.584 | 0.008 | - |
| Genetic effects | *Female genetic ancestry* | *0.270* | *0.022* | *more anubis ancestry in females → ↑ Pr(prox)* |
|  | **Male genetic ancestry** | **0.270** | **<0.001** | **more anubis ancestry in males → ↑ Pr(prox)** |
|  | **Assortative genetic ancestry index (*b*)** | **0.303** | **<0.001** | **females and males of similar genetic ancestry → ↑ Pr(prox)** |
|  | Female heterozygosity | -0.120 | 0.654 | - |
|  | Male heterozygosity | 0.042 | 0.745 | - |
|  | Genetic relatedness | -0.041 | 0.693 | - |
| Rank effects | **Female ordinal rank** | **-0.024** | **<0.001** | **higher ranking females → ↑ Pr(prox)** |
|  | **Male ordinal rank** | **-0.047** | **<0.001** | **higher ranking males → ↑ Pr(prox)** |
|  | *Female ordinal rank × male ordinal rank* | *0.001* | *0.011* | *females and males of similar rank → ↑ Pr(prox)* |
| Age effects | Female age ($a_u$) | -0.004 | 0.644 | - |
| | Female age transformed ($a_t$) | -0.028 | 0.916 | - |
| Demographic effects | Adult females in group | -0.018 | 0.073 | - |
|  | **Adult males in group** | **-0.036** | **0.005** | **more adult males in group → ↓ Pr(prox)** |
| Reproductive state effects | Reproductive state (pregnant = -1, lactating = +1) | 0.056 | 0.187 | - |
|  | Reproductive state (as above) × female genetic ancestry | 0.021 | 0.851 | - |
|  | Reproductive state (as above) × male genetic ancestry | -0.059 | 0.269 | - |
| Co-residency effects | **Pair co-residency** | **0.037** | **<0.001** | **longer co-residency → ↑ Pr(prox)** |
| Observer effects | **Observer effort** | **0.027** | **<0.001** | **greater observer effort → ↑ Pr(prox)** |

<sup>a</sup> All variables included in this table were fit as fixed effects in the multivariate logistic regression model. Male and female identity were fit as random effects.
<sup>b</sup> Predictor variables for which $p < 0.01$ are bolded and $p < 0.05$ are italicized.
<sup>c</sup> Pr(prox) = probability of proximity.

Similar patterns were observed for females, although the effect of female rank was weaker than for male rank. Grooming and proximity were more likely to occur if the female in a dyad had more anubis ancestry (grooming: β = 0.513, p < 0 .001, Table 1, Fig. 1c; proximity: β = 0.270, p = 0.022, Table 2, Fig. S2c) and was higher ranking (grooming: β = −0.027, p < 0.001, Table 1, Fig. 1d; proximity: β = −0.024, p < 0.001, Table 2, Fig. S2d). Female reproductive state, female age, transformed female age, and female heterozygosity did not significantly affect grooming or proximity behavior, nor did the interaction between female reproductive state and female genetic ancestry.

### Dyad-level characteristics: traits of both partners predict the propensity to affiliate with the opposite sex

In addition to individual-level effects, we found that the combined characteristics of the female and male in each dyad predicted the probability of grooming and proximity. First, affiliative interactions were assortative with respect to genetic ancestry: they were more likely to occur when both partners were of similar genetic ancestry (i.e., both anubis-like or both yellow-like) and less likely to occur if they were of different genetic ancestry (grooming: β = 0.646, p < 0.001, Table 1, Fig. 2; proximity: β = 0.303, p < 0.001, Table 2, Fig. S3). Overall, the probability of grooming and proximity was highest for pairs where both partners were anubis-like. Affiliative interactions were also assortative with respect to dominance rank: if both partners were high-ranking, the probability of affiliative interaction was higher than explained by the separate, additive effects of high male rank and high female rank alone (grooming: β = 0.003, p < 0.001, Table 1, Fig. 3; proximity: β = 0.001, p = 0.011, Table 2, Fig. S4). The effects of ancestry-based assortativity and rank-based assortativity are likely to be independent, as the assortative genetic ancestry index we used here is only weakly correlated with the product of male and female rank (the absolute value of Pearson’s product-moment correlation: *r* < 0.07, *P* < 3.7 × 10^−13^ for both grooming and proximity data sets). The correlation between rank and genetic ancestry is similarly weak within each sex (the absolute value of Pearson’s product-moment correlation: *r* < 0.03 for unique male rank-genetic ancestry combinations, *P* > 0.35 for both grooming and proximity; *r* < 0.08 for unique female rank-genetic ancestry combinations, *P* = 0.37 for grooming and *P* = 0.04 for proximity).

**Figure 2.**
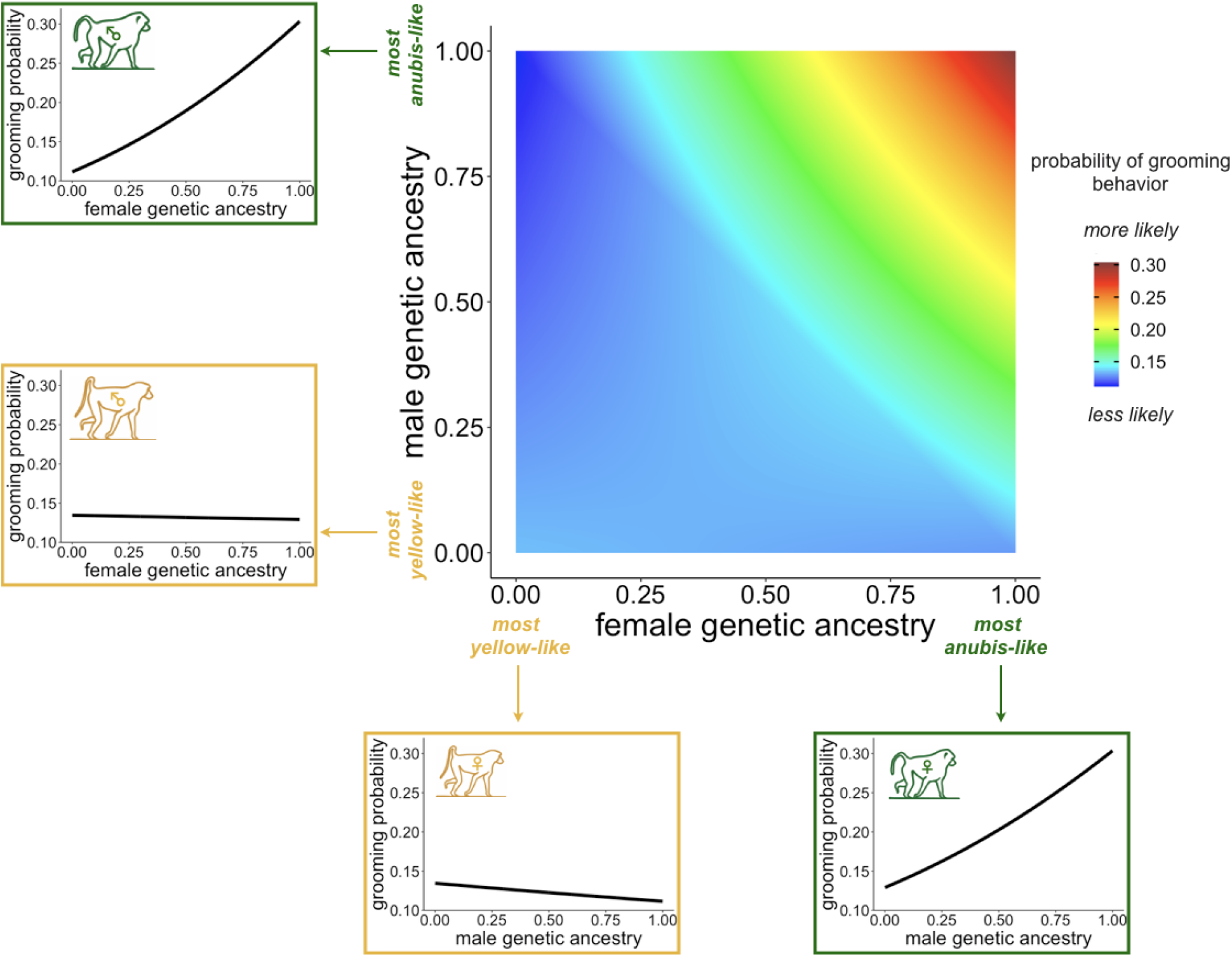
Combined genetic ancestry characteristics of females and males affect the probability of grooming. The central heatmap shows the probability of grooming behavior as a function of female genetic ancestry (x-axis) and male genetic ancestry (y-axis), based on model estimates assuming average values for all other covariates (see Supplementary Methods). Assortative affiliative behavior is reflected by increased probability of yellow-like females grooming with yellow-like males, relative to anubis-like males, and anubis-like females grooming with anubis-like males, relative to yellow-like males. The probability of grooming is highest for pairs where both partners are anubis-like. Line graphs surrounding the heatmap show model predictions for the probability of grooming behavior for males (left) and females (bottom) at the two extremes of genetic ancestry, as a function of the genetic ancestry of potential opposite-sex social partners. Baboon illustrations adapted from Alberts and Altmann (2001).

**Figure 3.**
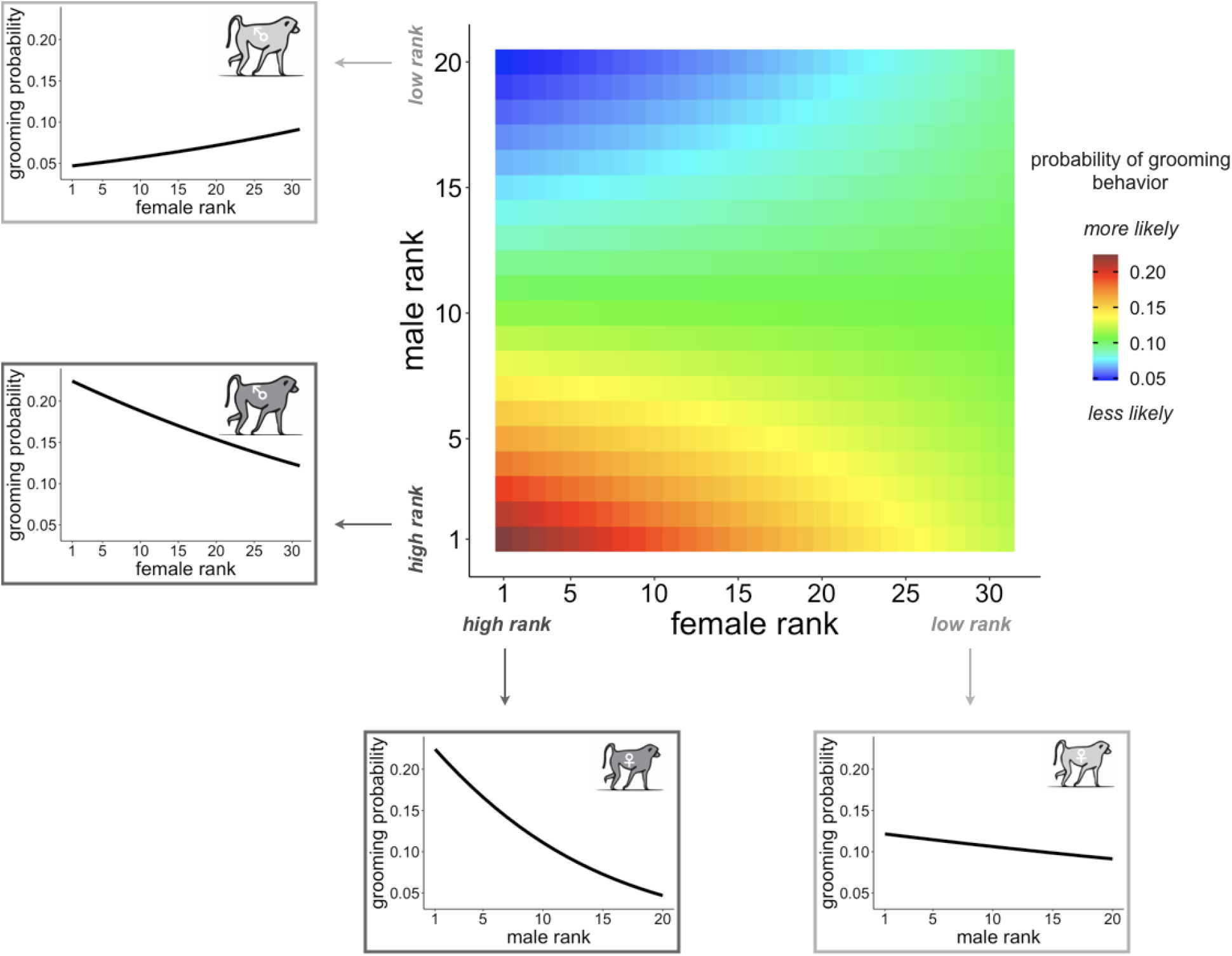
Combined rank characteristics of females and males affect the probability of grooming. The central heatmap shows the probability of grooming behavior as a function of female dominance rank (x-axis) and male dominance rank (y-axis), based on model estimates assuming average values for all other covariates (see Supplementary Methods). The probability of grooming is highest for pairs where both partners are high ranking. Line graphs surrounding the heatmap show model predictions for the probability of grooming behavior for males (left) and females (bottom) at the extremes of the rank distribution, as a function of the dominance rank of potential opposite-sex social partners.

Genetic relatedness did not predict either grooming or proximity behavior within dyads (grooming: Table 1; proximity: Table 2), in contrast to the effects of relatedness on mating behavior, where relatives are less likely to mate (consistent with inbreeding avoidance in baboons: Alberts & Altmann, 1995; Packer, 1979a; Tung et al., 2012). In other words, opposite-sex kin were neither more likely nor less likely to socially affiliate than opposite-sex nonkin. Additionally, male-female affiliation did not depend on the interaction between female reproductive state and male genetic ancestry (grooming: Table 1; proximity: Table 2).

### Group demography influences grooming and proximity behavior, and observer effort affects ascertainment of these behaviors

In addition to individual and dyadic-level effects, we found that aspects of group demography also influence male-female affiliative behavior. The probability of grooming was lower for all dyads when the social group contained more adult males (β = −0.030, p = 0.018, Table 1, Fig. S5a-b) and more adult females (β = −0.039, p < 0.001, Table 1, Fig. S6a-b). Similarly, the probability of proximity was lower for all dyads when the social group contained more adult males (β = −0.036, p = 0.005, Table 2, Fig. S5c-d), but not more adult females (β = −0.018, p = 0.073, Table 2, Fig. S6c-d).

Finally, the probability of grooming and proximity behavior was higher for all dyads the more days they were observed together in the same group (grooming: β = 0.047, p < 0.001, Table 1; proximity: β = 0.037, p < 0.001, Table 2). The probability of recording proximity behavior, but not grooming behavior, also increased with greater observer effort (grooming: β = 0.002, p = 0.425, Table 1; proximity: β = 0.027, p < 0.001, Table 2).

## DISCUSSION

Our results show that opposite-sex affiliative relationships are predicted by genetic ancestry in a natural baboon hybrid zone. Genetic ancestry effects are of particular interest because opposite-sex relationships must have a partial genetic basis in order to respond to natural selection. Additionally, genetic ancestry-associated differences provide *prima facie* evidence that this trait has evolved in the past. Specifically, in the Amboseli baboons, genetic ancestry acts alongside the effects of dominance rank and group demography to predict grooming and proximity behavior between adult males and adult females outside the mating context. These effects are not only detectable as a function of the individual characteristics of males and females, but also as a function of the properties of each opposite-sex pair. Although more anubis-like males and females were more likely to affiliate with the opposite sex regardless of their partner’s ancestry, pairs of anubis-like males and anubis-like females were the most likely to be observed grooming or in close proximity (Figs. 2, S3). Our findings thus suggest that the tendency to engage in opposite-sex affiliative behavior partially diverged during baboon evolution to differentiate yellow and anubis baboons. We note that while we tested for the effects of genetic ancestry in this study, not genotype *per se*, baboons in Amboseli inherit anubis ancestry from both maternal and paternal lines (Tung et al., 2008). Our data set also contains many multigeneration hybrids, such that genetic ancestry estimates vary continuously between mostly yellow to mostly anubis. Thus, the signature of genetic ancestry reported here likely arises from ancestry-associated differences in genotype, as opposed to ancestry-associated maternal or environmental effects on social preference.

These results add to previous evidence that male-female social bonds vary across baboon species (Baniel et al., 2016; Fischer et al., 2017; Goffe et al., 2016; Nguyen et al., 2009; Städele et al., 2019; Weingrill, 2000). For instance, male-female social relationships in chacma baboons are short-lived and occur primarily when females have dependent infants, whereas in Guinea baboons, close male-female social relationships commonly last for several years (Baniel et al., 2016; Fischer et al., 2017; Goffe et al., 2016). Yellow and anubis baboons, which have similar social organization and mating systems (multi-male, multi-female groups with female-biased dispersal and polygynandry), are thought to fall between these two extremes, such that male-female relationships can sometimes, although not always, be long-lasting (Nguyen et al., 2009; Smuts, 1985; Städele et al., 2019). The identification of genetic ancestry effects in this study thus suggests that subtle differences in the nature of opposite-sex social relationships can evolve even between species that are otherwise quite similar in their social systems and behavioral repertoires. Identifying the molecular and neurochemical substrates for these differences, including whether they are shared with other taxa, is a fascinating topic for future work that could be facilitated by studies within natural hybrid zones.

Several lines of evidence also support the relevance of genetic ancestry effects on opposite-sex affiliative behavior to current variation in fitness. First, opposite-sex social relationships predict longevity in the Amboseli baboons (Archie et al., 2014; Campos et al., 2020), and longevity is an important contributor to lifetime reproductive success in both male and female baboons, as well as in other long-lived vertebrates (Alberts et al., 2006; Clutton-Brock, 1988; Lawler, 2007; McDonald, 1993; McLean et al., 2019; Newton, 1989; Wroblewski et al., 2009). Second, male-female social bonds can also lead to other reproductive gains, including offspring care that may improve survival (Anderson, 1992; Buchan et al., 2003; Busse & Hamilton, 1981; Huchard et al., 2013; Moscovice et al., 2009; Nguyen et al., 2009; Silk et al., 2020). Indeed, our findings that affiliative behavior was less common for any given dyad in large groups, and that both male and female rank predicted social interactions, suggest that male-female social bonds are an important and limited social resource for both sexes (Archie et al., 2014; Baniel et al., 2016; Haunhorst et al., 2019; Lemasson et al., 2008; Palombit et al., 2001; Seyfarth, 1976; Städele et al., 2019). This interpretation agrees with reports in chacma baboons that pregnant and lactating females direct aggression towards cycling females that are mate-guarded by and copulate with a shared male social partner (Baniel et al., 2018). Together with ancestry-related differences in affiliative behavior, our observations indicate that opposite-sex affiliative behavior has not only evolved in baboons in the past, but may also be the target of selection in the Amboseli population today.

The long-term ramifications of our findings for the stability or resolution of the hybrid zone remain somewhat unclear. If strong opposite-sex social bonds are fitness-enhancing, more anubis-like ancestry should be favored in Amboseli. However, assortative social preferences (both in mating and non-mating contexts) can also act as a barrier to admixture and could reduce the rate of anubis expansion. Along with previous findings in Amboseli (Charpentier et al., 2008; Franz et al., 2015; Tung et al., 2012), our results thus suggest that genetic ancestry is associated with a range of selectively relevant behavioral and life-history traits that do not universally point towards either anubis range expansion or to behaviorally-mediated reproductive isolation. Furthermore, any effect of genetic ancestry in baboons must necessarily be filtered through the effect of dominance rank, which is the most robust predictor of male mating behavior and, based on the results of this study, also a major contributor to opposite-sex affiliative behavior (Tung et al., 2012). Finally, the effect of assortativity necessarily depends on the characteristics of available social partners. The interplay between genetic ancestry effects on an individual-level and at the dyadic-level will therefore be dynamic across populations and over time. The complexity of these co-acting factors may help explain why yellow baboons and anubis baboons remain phenotypically and genetically distinct, even though genomic analyses indicate repeated bouts of gene flow between yellow baboons and anubis baboons over hundreds to thousands of generations (Rogers et al., 2019; Wall et al., 2016). More broadly, our results suggest that simple behavioral barriers to admixture, such as wing pattern-based mate choice in butterflies or vibrational-based courtship signals in treehoppers (Jiggins et al., 2001; Rodríguez et al., 2004), are unlikely to occur in socially complex animals like baboons.

Finally, our study suggests both parallels and differences between opposite-sex social interactions within versus outside the context of mating. Several predictors of opposite-sex affiliation in this study overlap with predictors of mating behavior, including increased probability of both mating and social affiliation for higher-ranking and more anubis-like males, and assortativity based on both genetic ancestry and dominance rank (Tung et al., 2012). However, effects of female age and kinship were weak or undetectable in our analysis of male-female social bonds outside of the mating context. These results contrast with our previous result that male baboons are less likely to mate with females in older age classes (Tung et al., 2012), and with multiple lines of evidence that baboons and other primates avoid mating with relatives (Alberts & Altmann, 1995; Godoy et al., 2016; Packer, 1979a; Tung et al., 2012; Walker et al., 2017; Widdig et al., 2017). Because opposite-sex social affiliation is generally not a strong predictor of future mating events (although it may reflect a past history of mating: Huchard et al., 2010; Moscovice et al., 2010; Nguyen et al., 2009; Silk et al., 2020; Städele et al., 2019), differences in male-female behavior in mating versus non-mating contexts may arise because the benefits of opposite-sex social bonds differ from the benefits of mating. Alternatively or additionally, female choice and female-female competition may be more important in predicting grooming and proximity than mating behavior. Whereas only one or a few females experience estrus at any given time (Bercovitch, 1983; Bulger, 1993; Levy et al., 2020), all adult females are available as, and may actively be searching out, grooming partners. Notably, grooming relationships in baboons are more often initiated and maintained by females than by males, whereas males primarily absorb the costs of mate-guarding in a mating context (Alberts et al., 1996; Nguyen et al., 2009; Packer, 1979b; Palombit et al., 1997; but see Weyher et al., 2014). Thus, sexual and social preferences, and the extent to which they are expressed by males versus females, are likely to vary across different types of opposite-sex relationships—a distinction reminiscent of differences between social monogamy and genetic monogamy in pair-bonded birds and mammals (Carter & Perkeybile, 2018; Gowaty, 1996).

## ACKNOWLEDGEMENTS

We gratefully acknowledge the support of the National Science Foundation and the National Institutes of Health for the majority of the data represented here, currently through NSF IOS 1456832, NIH R01AG053308, R01AG053330, R01HD088558, and P01AG031719. We also thank the North Carolina Biotechnology Center for support for high-performance computing resources (2016-IDG-1013). A.S.F. was supported by NSF GRFP (DGE #1644868) and NIH T32GM007754; E.M.M was supported by NSF IOS 1501971. We thank Duke University, Princeton University, and the University of Notre Dame for financial and logistical support. In Kenya, we thank the Kenya Wildlife Service, University of Nairobi, Institute of Primate Research, National Museums of Kenya, the National Environment Management Authority (NEMA), and the National Council for Science, Technology, and Innovation (NACOSTI). We also thank the members of the Amboseli-Longido pastoralist communities, the Enduimet Wildlife Management Area, Ker & Downey Safaris, Air Kenya, and Safarilink for their cooperation and assistance in the field. Particular thanks go to the Amboseli Baboon Research Project field team (R.S. Mututua, S. Sayialel, J.K. Warutere, I.L. Siodi, G. Marinka, B. Oyath) and camp staff, and to T. Wango and V. Oudu for their untiring assistance in Nairobi, and to Jeanne Altmann for her fundamental contributions to the Amboseli baboon research. The baboon project database, Babase, was designed and programmed by K. Pinc and is expertly managed by N.H. Learn and J.B. Gordon. Finally, we thank current and previous members of the Alberts and Tung labs for their helpful feedback. For a complete set of acknowledgments of funding sources, logistical assistance, and data collection and management, please visit http://amboselibaboons.nd.edu/acknowledgements/. Any opinions, findings, and conclusions expressed in this material are those of the authors and do not necessarily reflect the views of our funding bodies.

## DATA STATEMENT

Data on grooming behavior, proximity behavior, and all predictor variables tested in our models will be available on Dryad upon acceptance.

