## Supplementary Material for "Genetic ancestry predicts male-female affiliation in a natural baboon hybrid zone"

### Supplementary Methods

*Estimating observer effort.* All baboon study groups are observed by the same number of field observers for roughly the same amount of time, regardless of group size. As a consequence, the individual-level density of our grooming and proximity data varies by social group (i.e., there are differences among groups in observer effort per individual baboon). To account for these differences, we estimated observer effort using focal animal sampling data from adult females. Each focal sample typically consists of 10 point samples, collected once per minute, for 10 minutes (Alberts et al., 2020; Archie et al., 2014). We summed the total number of point samples collected for adult females per social group across each female's two-month interval and then divided this value by the total number of adult females present in the social group during that same time period. We used this value as a measure of observer effort and included it as a fixed effect covariate in our models.

*Filtering of grooming and proximity data.* In the main text, we report results for grooming and proximity behavior using two-month time intervals. Initially, we attempted to measure grooming and proximity behavior on a monthly basis. However, on a monthly basis, grooming and proximity were not observed in 93.2% and 87.2% of rows (n=62,195 total rows), respectively. While some of these "0" value rows reflect a true absence of male-female interactions, observations in a single month may be too sparse to accurately reflect whether a given male-female dyad interacted. Indeed, even after omitting social group-month combinations with low observer effort (i.e., less than an average of 20 point samples per female per social group-month), grooming and proximity were still not observed in 92.1% and 83.9% of the data set, respectively. Thus, we concluded that our resolution of grooming and proximity behavior was too coarse to measure on a monthly basis.

To address this concern, we expanded our time window from one month to two months and excluded any male in a female's two-month interval if he was only present for one of the two months. This decision reduced the percentage of rows where grooming and proximity did not occur (87.9% and 76.9% of rows, respectively, n=26,245 total rows). Because we were most interested in the predictors of opposite-sex affiliative behavior, conditioned on it actually occurring, we also filtered out any two-month interval where the focal female did not interact with any male social partner for the entire interval (31.7% and 17.1% of intervals for the grooming and proximity data sets, respectively). We performed this filtering separately for the grooming and proximity data sets, thus producing two separate data sets for downstream analysis. We also excluded all data for females and males who were observed for less than 8 months. If these filters resulted in no observations of grooming or proximity for a given female interval, we also removed that interval (and repeated this procedure until all filtering criteria were met). Finally, we did not consider two-month intervals in which females transitioned between reproductive states (pregnant or lactating) and two-month intervals in which there were fewer than two adult males available for a female to interact with.

*Encoding female reproductive state.* In the main text, we report results for grooming and proximity behavior from models where female reproductive state was coded as -1 for pregnancy and 1 for lactation. This deviates from the usual arbitrary coding of binary states as 0/1. We made this decision because we found that, with the 0/1 encoding, our beta and p-value estimates for some model parameters depended on whether we assigned pregnancy to the 0 state versus lactation. Using the -1/1 alternative encoding eliminated this issue, produced consistent results across R packages (*glmmTMB*, version 1.0.1 (Brooks et al., 2017) versus *lme4,* version 1.1.23 (Bates et al., 2015)), and also qualitatively matched our estimates using a fixed effects-only model (using the function *glm* from base R). Hence, we report the results using the -1/1 encoding in the main text.

*Visualizing model effects.* For Figures 1b, 1d, 2, 3, and Supplementary Figures S2b, S2d, S3, and S4, we plotted the probability of grooming and proximity behavior as a function of a varying predictor of interest and model estimates assuming average values for all other covariates. For example, to calculate the probability of grooming and proximity as a function of male dominance rank, we modeled an "average" male (apart from his rank) interacting with an "average" female in an "average" demographic environment. We then estimated how variation in male rank is predicted to affect the probability of grooming or proximity using R's *predict* function. For predictors that reflect the combined characteristics of the male-female pair, we used the same approach, but extended it to vary the characteristic of interest for both the male and the female, while holding the other predictor variables constant at their average value.

Supplementary Figures and Tables

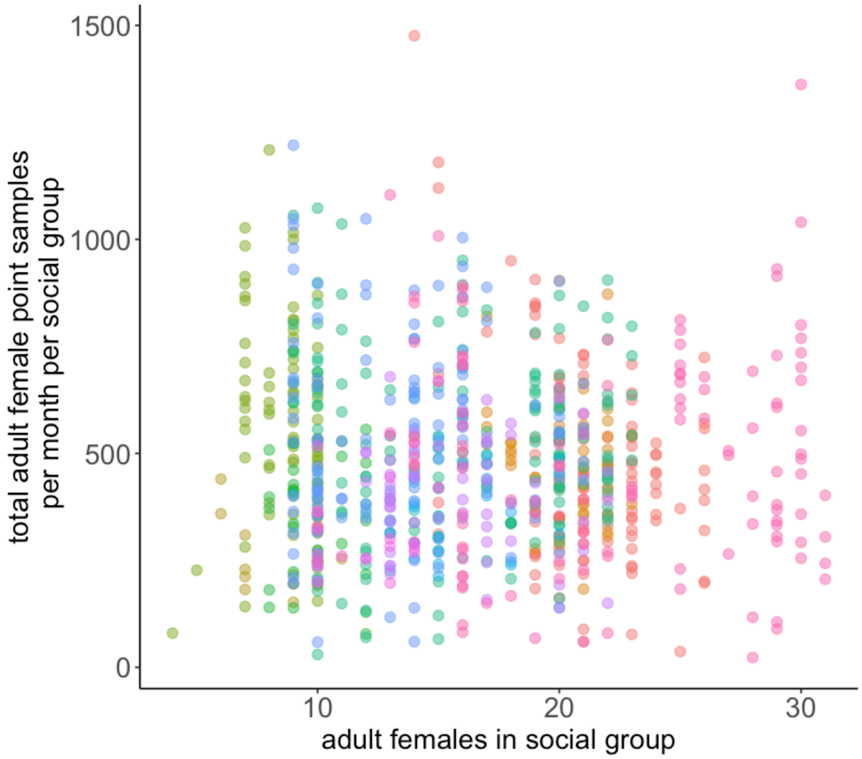


**Figure S1. The total effort invested in behavioral observations is consistent across study groups of different sizes.** Each dot represents a unique social group-month combination, colored by social group (n=12 social groups, n=812 total group-months from the initial, monthly-based data set). There is no relationship between the number of adult females present in a social group and the total number of point samples recorded for adult females per social group (10 point samples are collected per focal sample, if the sample is complete; linear model estimate for the effect of number of adult females on total number of point samples: $\beta = -1.62$, $p = 0.221$). The result is that observer effort per individual baboon varies with group size, and the probability of observing true grooming or proximity events is smaller for large groups than for small groups. To address this problem, we included a measure of observer effort (total adult female point samples per social group / total adult females present in social group) in our models.

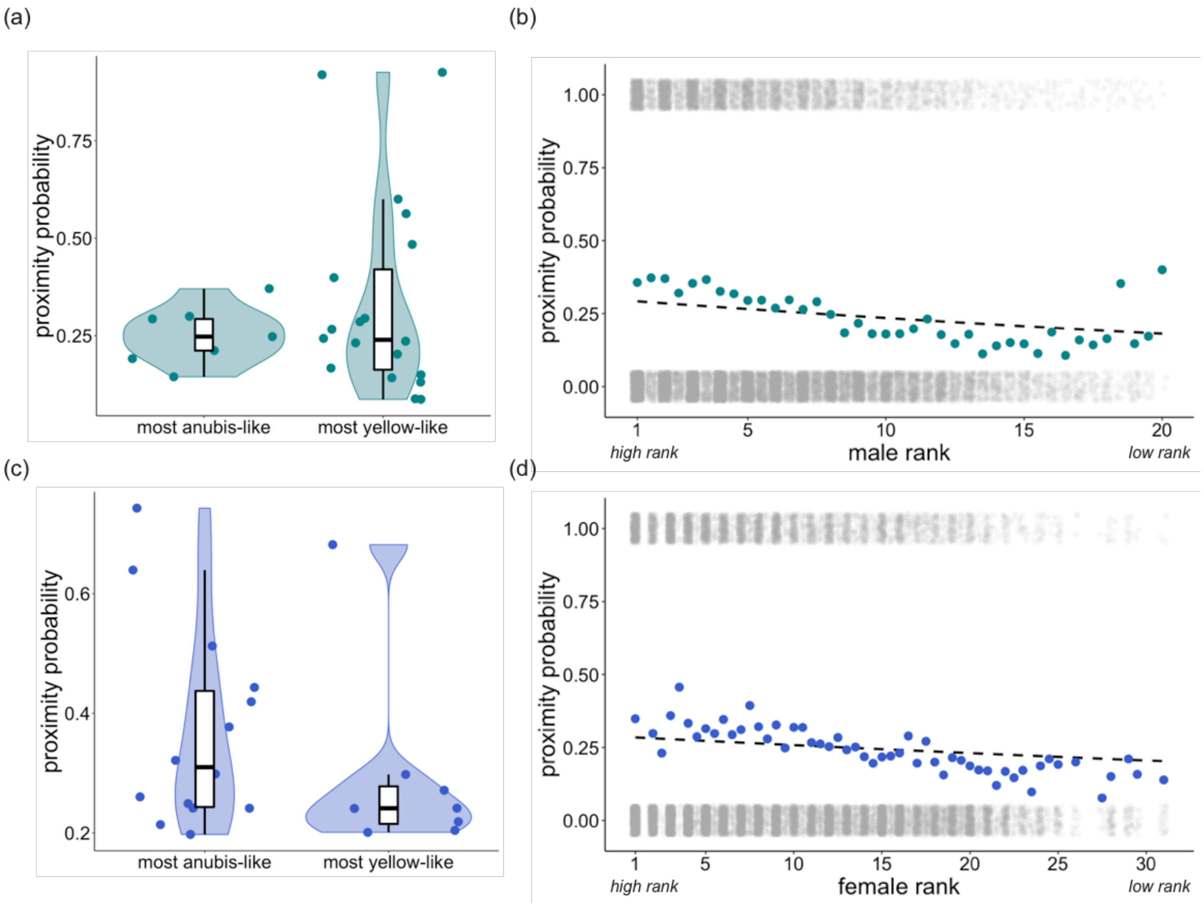


**Figure S2. Genetic ancestry and dominance rank predict the tendency to be in proximity to an opposite-sex partner. (a)** The probability of being in proximity among co-resident opposite-sex pairs, per two-month interval, for the most anubis-like males (above the 90th percentile for male genetic ancestry in the data set, > 81.5% anubis ancestry, n=9 males) and the most yellow-like males (below the 10th percentile for male genetic ancestry in the data set, < 4.8% anubis ancestry, n=20 males). Probabilities were calculated from the data without adjustment for other covariates (note that, although not visually apparent in the 1/0 proximity data as summarized here, more anubis-like males are more likely to be observed in proximity with female partners in the full model: $\beta = 0.270$, $p < 0.001$, Table 2). **(b)** The probability of being in proximity among co-resident opposite-sex pairs, per two-month interval, as a function of male dominance rank. Colored dots show probabilities based on counts of proximity occurrences, without adjustment for other covariates (as in (a)), and the dashed line shows the predicted relationship based on model estimates, assuming average values for all other covariates (see Supplementary Methods). Grey dots show the presence (y=1) or absence (y=0) of proximity behavior for all 21,130 female-male pair-interval combinations, as a function of male dominance rank (dots are jittered vertically for visibility). Non-integer values correspond to individuals that changed ranks during a two-month interval in the data set. **(c)** As in (a), for the most anubis-like females (above the 90th percentile for female genetic ancestry in the data set, > 74.1% anubis ancestry, n=14 females) and the most yellow-like females (below the 10th percentile for female genetic ancestry in the data set, < 3.2% anubis ancestry, n=8 females). **(d)** As in (b), with the probability of being in proximity shown as a function of female dominance rank.

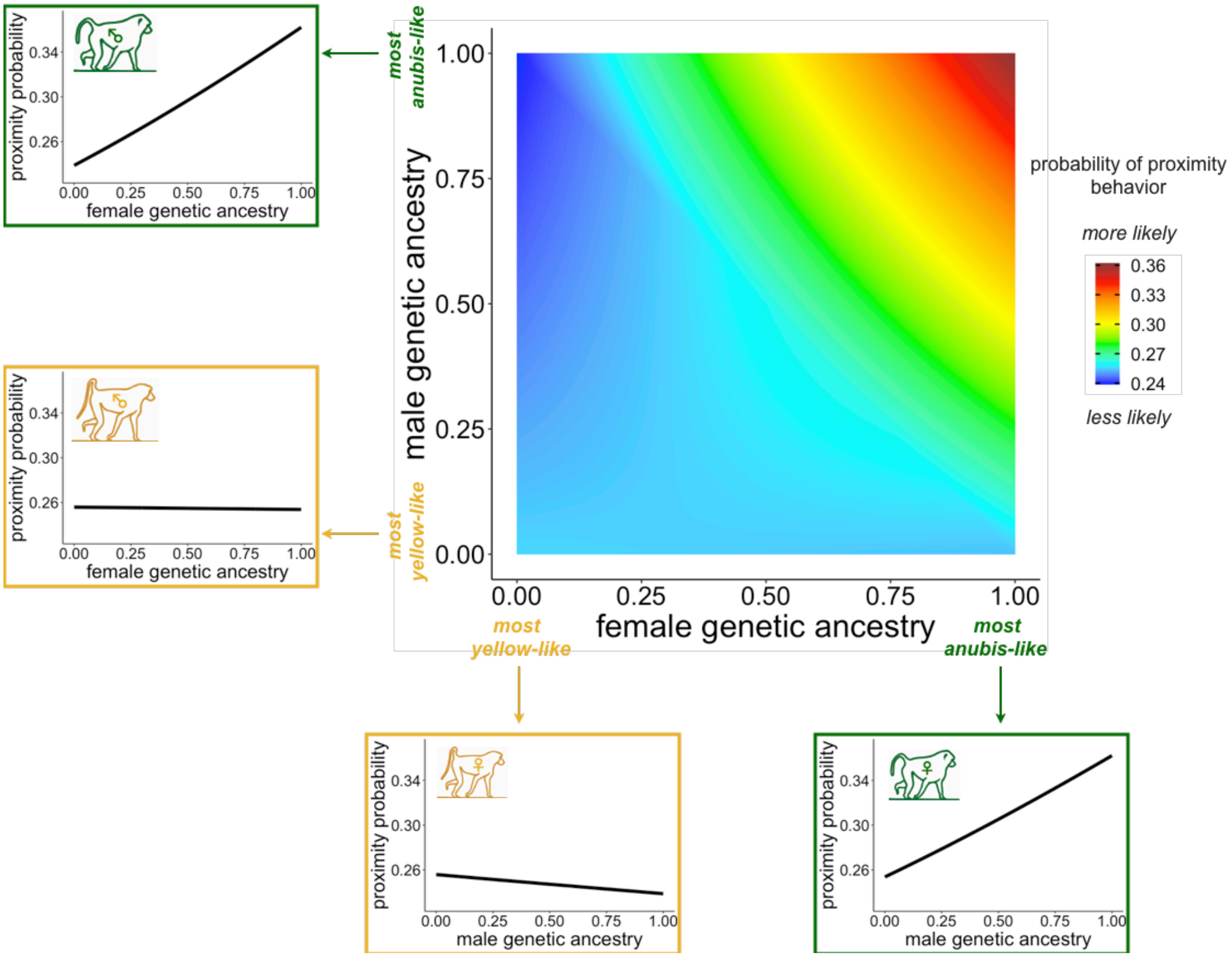


**Figure S3. Combined genetic ancestry characteristics of females and males affect the probability of being in proximity.** The central heatmap shows the probability of proximity behavior as a function of female genetic ancestry (x-axis) and male genetic ancestry (y-axis), based on model estimates assuming average values for all other covariates (see Supplementary Methods). Assortative affiliative behavior is reflected by increased probability of yellow-like females being in proximity with yellow-like males, relative to anubis-like males, and anubis-like females being in proximity with anubis-like males, relative to yellow-like males. The probability of being in proximity is highest for pairs where both partners are anubis-like. Line graphs surrounding the heatmap show model predictions for the probability of proximity behavior for males (left) and females (bottom) at the two extremes of genetic ancestry, as a function of the genetic ancestry of potential opposite-sex social partners. Baboon illustrations adapted from Alberts and Altmann (2001).

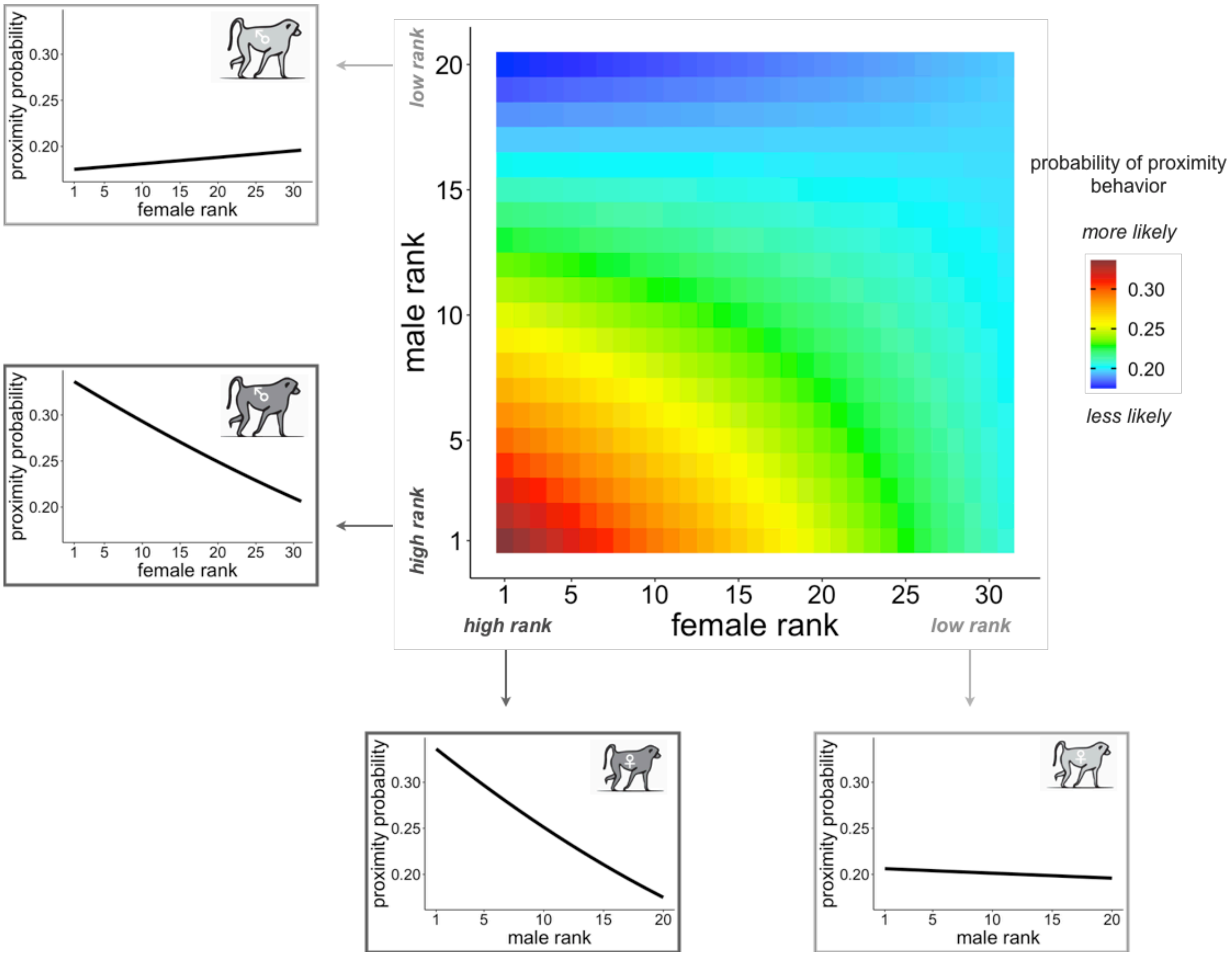


**Figure S4. Combined rank characteristics of females and males affect the probability of being in proximity.** The central heatmap shows the probability of proximity behavior as a function of female dominance rank (x-axis) and male dominance rank (y-axis), based on model estimates assuming average values for all other covariates (see Supplementary Methods). The probability of being in proximity is highest for pairs where both partners are high ranking. Line graphs surrounding the heatmap show model predictions for the probability of proximity behavior for males (left) and females (bottom) at the extremes of the rank distribution, as a function of the dominance rank of potential opposite-sex social partners.

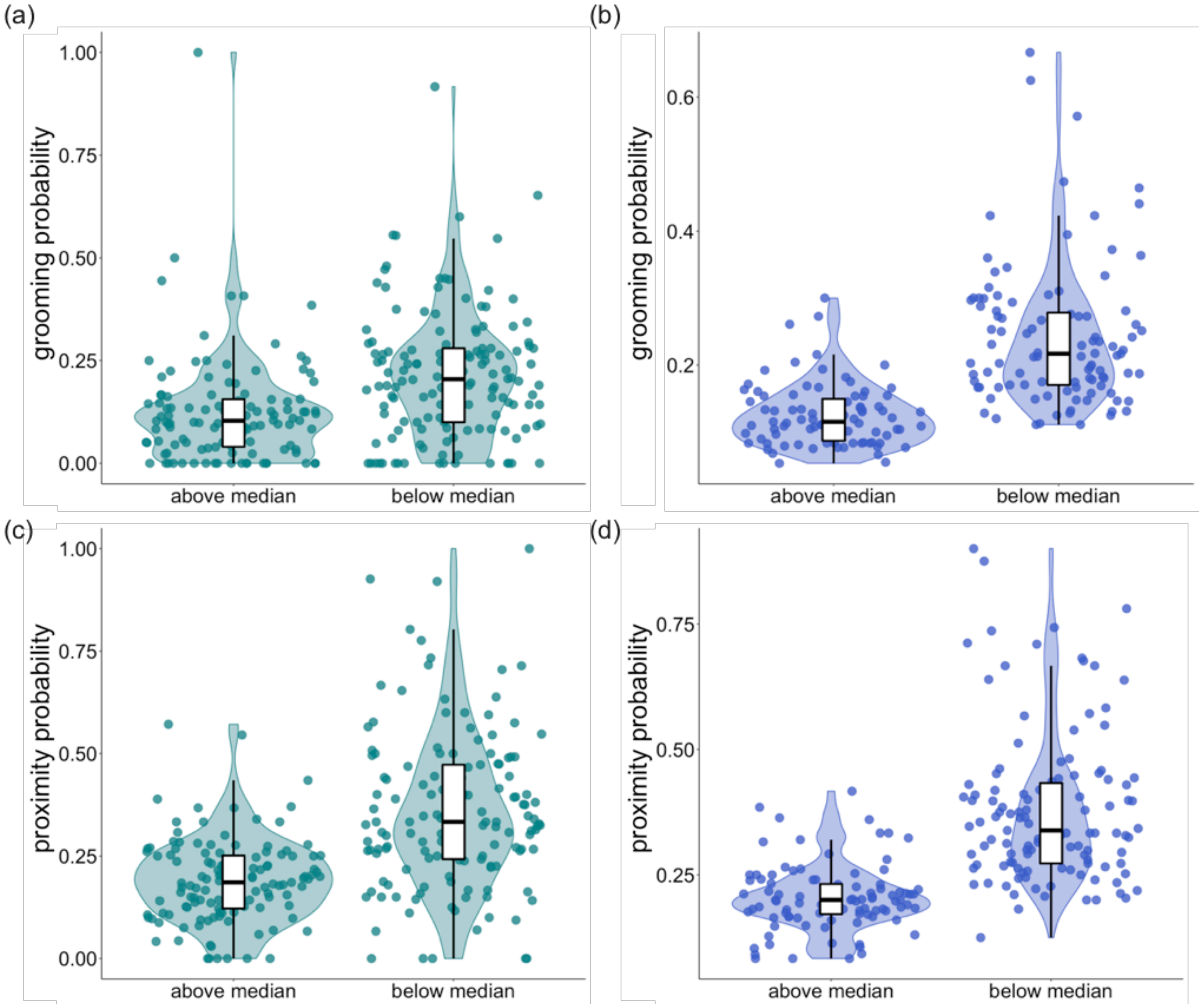


**Figure S5. The number of adult males present in a social group influences grooming and proximity behavior. (a-b)** The probability of grooming among co-resident opposite-sex pairs, per two-month interval, for each male (**a**) and female (**b**) in social groups with greater or less than the median number of co-resident males in the sample (n = 12 males). Probabilities were calculated from the data without adjustment for other covariates. **(c-d)** As in (a-b), for proximity behavior (median = 11.5 males).

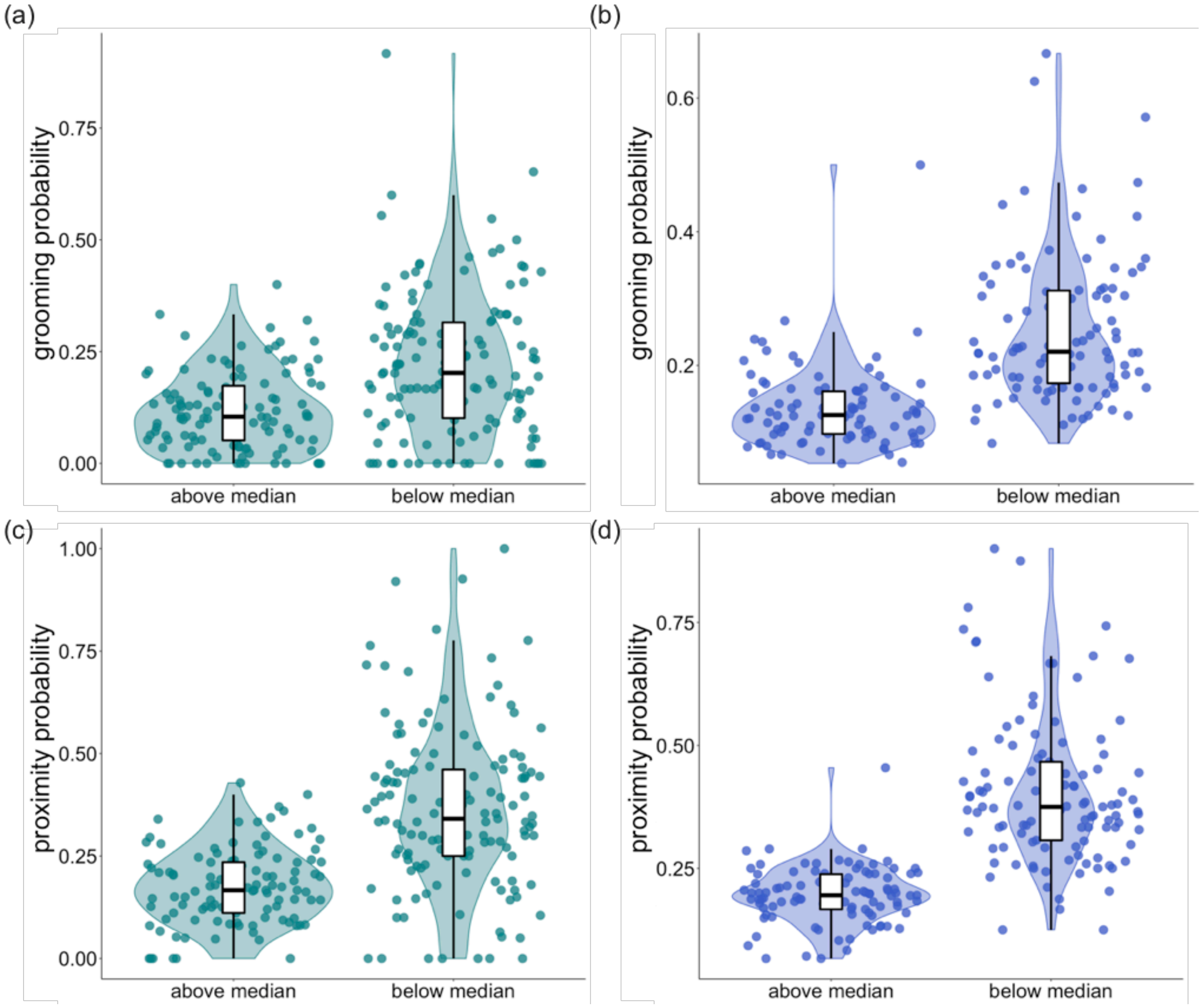


**Figure S6. The number of adult females present in a social group influences grooming but not proximity behavior. (a-b)** The probability of grooming among co-resident opposite-sex pairs, per two-month interval, for each male **(a)** and female **(b)** in social groups with greater or less than the median number of co-resident females in the sample (n = 20 females). Probabilities were calculated from the data without adjustment for other covariates. **(c-d)** As in (a-b), for proximity behavior (median = 20 females). Note that in the full model, the apparent difference observable for proximity is not significant after taking into account other model covariates and random effects ($\beta$ = -0.018, p = 0.073, Table 2).

**Supplementary Table 1.** Summary of social groups, study subjects, and observation periods.

| Group ID | Females[a] | Males[a] | Months in grooming data set[b] | | Months in proximity data set[b] | | Observation period[c] | Observation months[d] |
|---|---|---|---|---|---|---|---|---|
| | | | Per female | Per male | Per female | Per male | | |
| 1.1 | 33 | 48 | 21.88 ± 13.60 | 28.02 ± 23.40 | 30.06 ± 18.76 | 28.04 ± 23.38 | November 1999-September 2010 | 119 |
| 1.11 | 17 | 23 | 14.53 ± 8.19 | 20.52 ± 18.07 | 21.25 ± 6.81 | 21.09 ± 18.54 | May 2011-December 2015 | 56 |
| 1.12 | 5 | 9 | 10.40 ± 1.67 | 9.22 ± 5.83 | 12.00 ± 2.00 | 9.22 ± 5.83 | May 2011-November 2012 | 19 |
| 1.21 | 12 | 24 | 17.78 ± 10.74 | 13.25 ± 13.60 | 19.27 ± 12.78 | 13.46 ± 14.04 | November 1999-September 2008 | 72 |
| 1.211 | 8 | 13 | 16.50 ± 7.07 | 19.31 ± 10.55 | 19.75 ± 7.59 | 19.38 ± 10.52 | August 2012-December 2015 | 41 |
| 1.22 | 25 | 35 | 31.84 ± 18.97 | 29.80 ± 26.77 | 38.08 ± 21.60 | 30.29 ± 27.19 | November 1999-November 2012 | 145 |
| 1.221 | 8 | 9 | 6.33 ± 2.34 | 9.56 ± 5.10 | 11.25 ± 1.83 | 11.56 ± 4.82 | June 2014-December 2015 | 19 |
| 1.222 | 11 | 9 | 7.27 ± 2.57 | 12.22 ± 7.03 | 8.91 ± 2.74 | 12.89 ± 7.37 | June 2014-December 2015 | 19 |
| 2.1 | 20 | 44 | 27.50 ± 20.01 | 24.23 ± 19.73 | 32.50 ± 24.96 | 24.68 ± 19.89 | November 1999-March 2011 | 124 |
| 2.11 | 12 | 21 | 15.80 ± 7.08 | 17.67 ± 13.44 | 20.17 ± 5.75 | 18.14 ± 13.64 | June 2011-December 2015 | 55 |
| 2.12 | 8 | 10 | 6.33 ± 1.97 | 9.40 ± 6.98 | 8.75 ± 3.01 | 9.90 ± 7.36 | June 2011-November 2012 | 18 |
| 2.2 | 31 | 48 | 25.42 ± 13.54 | 29.27 ± 26.48 | 31.19 ± 15.62 | 29.52 ± 26.66 | November 1999-September 2011 | 121 |

[a] Number of unique individuals included in this data set, per unique social group. Individuals may be represented in more than one social group because of social group fissions and fusions and/or secondary dispersal by males.
[b] Mean ± s.d. Due to our filtering criteria, individuals must be in the data sets for at least 8 months but could be counted as members of different social groups.
[c] The observation starting and ending month and year for each social group across both grooming and proximity data sets. Some observation periods span time periods in which behavioral monitoring was inconsistent or when social groups were too unstable (i.e., social groups were fissioning or fusing) to unambiguously determine an individual's group membership. For our analysis, we excluded these time periods as well as the 2009 hydrological year (November 1st, 2008-October 31st, 2009). We also excluded months when the number of adult males in the social group was less than two.
[d] The number of unique observation months for each social group included across the grooming and proximity data sets.

**Supplementary Table 2.** Pearson's product-moment correlation (*r*) between predictor variables in the final grooming data set.

| | Genetic ancestry | | Heterozygosity | | | Dominance rank | | | Adults in social group | | | | |
|---|---|---|---|---|---|---|---|---|---|---|---|---|---|
| | Male | Assortative genetic ancestry index | Female | Male | Genetic relatedness | Female | Male | Female age | Female | Male | Reproductive state | Pair co-residency | Observer effort |
| Genetic ancestry | | | | | | | | | | | | | |
| Female | **0.098** | **-0.391** | **0.282** | **0.070** | **-0.086** | **0.027** | **0.028** | -0.009 | **0.189** | **0.110** | **0.020** | **-0.053** | **-0.054** |
| Male | | **-0.612** | **0.048** | **0.292** | **-0.158** | **0.047** | *-0.016* | **0.036** | **0.115** | **0.108** | -0.002 | **0.042** | **-0.062** |
| Assortative genetic ancestry index | | | **-0.131** | **-0.159** | **0.244** | **-0.035** | **-0.069** | -0.001 | **-0.090** | **-0.082** | 0.001 | **-0.044** | *0.017* |
| Heterozygosity | | | | | | | | | | | | | |
| Female | | | | **0.044** | **-0.091** | **0.185** | **0.067** | *-0.015* | **0.205** | **0.151** | 0.005 | **-0.034** | **-0.152** |
| Male | | | | | **-0.086** | **0.020** | 0.010 | 0.009 | **0.056** | **0.077** | -0.013 | 0.000 | **-0.042** |
| Genetic relatedness | | | | | | 0.011 | **-0.045** | -0.005 | **-0.030** | **-0.029** | -0.002 | **0.033** | 0.010 |
| Dominance rank | | | | | | | | | | | | | |
| Female | | | | | | | **0.167** | -0.007 | **0.396** | **0.335** | **0.026** | **0.039** | **-0.255** |
| Male | | | | | | | | 0.014 | **0.399** | **0.505** | **-0.028** | **-0.033** | **-0.220** |
| Female age | | | | | | | | | **0.025** | **0.021** | **0.068** | **0.076** | **-0.064** |
| Adults in social group | | | | | | | | | | | | | |
| Female | | | | | | | | | | **0.804** | *0.018* | **-0.029** | **-0.614** |
| Male | | | | | | | | | | | **-0.050** | **0.039** | **-0.436** |
| Reproductive state | | | | | | | | | | | | **-0.027** | *0.017* |
| Pair co-residency | | | | | | | | | | | | | **0.026** |

Predictor variables for which $p < 0.01$ are bolded and $p < 0.05$ are italicized.

**Supplementary Table 3.** Pearson's product-moment correlation (*r*) between predictor variables in the final proximity data set.

| | Genetic ancestry | | Heterozygosity | | | Dominance rank | | | Adults in social group | | | | |
|---|---|---|---|---|---|---|---|---|---|---|---|---|---|
| | Male | Assortative genetic ancestry index | Female | Male | Genetic relatedness | Female | Male | Female age | Female | Male | Reproductive state | Pair co-residency | Observer effort |
| Genetic ancestry | | | | | | | | | | | | | |
| Female | **0.095** | **-0.390** | **0.253** | **0.071** | **-0.086** | -0.008 | **0.021** | 0.001 | **0.160** | **0.098** | 0.011 | **-0.032** | **-0.040** |
| Male | | **-0.655** | **0.050** | **0.272** | **-0.164** | **0.032** | **-0.021** | **0.039** | **0.102** | **0.090** | -0.001 | **0.053** | **-0.052** |
| Assortative genetic ancestry index | | | **-0.124** | **-0.163** | **0.237** | *-0.016* | **-0.056** | -0.008 | **-0.068** | **-0.060** | 0.001 | **-0.056** | 0.011 |
| Heterozygosity | | | | | | | | | | | | | |
| Female | | | | **0.045** | **-0.086** | **0.190** | **0.045** | **-0.027** | **0.189** | **0.116** | -0.004 | **-0.028** | **-0.148** |
| Male | | | | | **-0.087** | *0.016* | 0.000 | *0.015* | **0.056** | **0.074** | -0.009 | 0.003 | **-0.045** |
| Genetic relatedness | | | | | | **0.020** | **-0.041** | 0.002 | *-0.017* | *-0.015* | 0.002 | **0.036** | 0.001 |
| Dominance rank | | | | | | | | | | | | | |
| Female | | | | | | | **0.150** | 0.013 | **0.378** | **0.304** | **0.066** | **0.029** | **-0.253** |
| Male | | | | | | | | **0.050** | **0.375** | **0.491** | **-0.026** | **-0.037** | **-0.222** |
| Female age | | | | | | | | | **0.089** | **0.095** | **0.070** | **0.077** | **-0.101** |
| Adults in social group | | | | | | | | | | | | | |
| Female | | | | | | | | | | **0.781** | *0.016* | **-0.028** | **-0.637** |
| Male | | | | | | | | | | | **-0.051** | **0.035** | **-0.451** |
| Reproductive state | | | | | | | | | | | | *-0.016* | 0.002 |
| Pair co-residency | | | | | | | | | | | | | **0.022** |

Predictor variables for which $p < 0.01$ are bolded and $p < 0.05$ are italicized.
